# Generalizable Cysteine Quantification in Pea Cultivars from SERS Spectra Using AI

**DOI:** 10.64898/2026.03.20.713189

**Authors:** Elham Gorgannejad, Qian Liu, Catherine Rui Jin Findlay, Mohammad Nadimi, Alex Chun-Te Ko, Pankaj Bhowmik, Jitendra Paliwal

## Abstract

Rapid quantification of sulfur-containing amino acids, particularly cysteine, in legumes is critical for assessing nutritional quality, supporting breeding program screening, and ensuring consistency in quality control processes. However, conventional methods, such as high-performance liquid chromatography (HPLC), are time-consuming and resource-intensive for high-throughput applications. This study evaluated artificial intelligence models for predicting cysteine concentration from surface-enhanced Raman spectroscopy (SERS) spectra of pea extracts. SERS spectra were acquired from 20 cultivars grown at three geographically distinct locations, with HPLC-measured cysteine concentrations as a ground truth reference. Linear regression, partial least squares regression, support vector regression, random forest regression, and a one-dimensional convolutional neural network (1D-CNN) were compared using within-cultivar splits and leave-one-cultivar-out (LOCO) evaluation. The 1D-CNN achieved RMSE 0.008 g/100 g within cultivars and maintained performance under LOCO, while other models showed limited generalization. Shapley Additive Explanations highlighted informative bands in the 630–760 cm^−1^ range, and noise modeling optimized scan-count selection.

## 1. Introduction

Legumes are an important source of plant-based protein in human diets (Lisciani et al., 2024; Samal et al., 2023). Their seeds contain approximately 20–45% protein on a dry weight basis, depending on species and cultivar (Maphosa et al., 2017). This is higher than the protein content of most widely consumed plant-based foods, including cereals (7–15%) and vegetables (1–5%) (Boye et al., 2010). Despite this advantage, legume protein quality is constrained by low levels of the sulfur-containing amino acids (SCAAs), cysteine and methionine (Iqbal et al., 2006). Together, these two amino acids can at times constitute the primary limiting essential and semi-essential amino acids determining protein quality. Peas, beans, and lentils typically contain only 12–18 mg/g protein of cysteine plus methionine, which is below the 22–25 mg/g protein recommended in the Food and Agriculture Organization/World Health Organization (FAO/WHO) amino acid reference pattern for high-quality dietary protein (WHO/FAO/UNU, 2025). SCAAs levels are influenced by cultivar genetics and environmental conditions, such as soil type, climate, and agronomic practices, as well as by genotype-by-environment (G×E) interactions (Gerrano et al., 2022). High-throughput, cultivar-robust quantification methods are important for the development of reliable, routine identification of high-SCAA genotypes and for quality control in commercial legume protein ingredients. Conventional analytical methods, such as high-performance liquid chromatography (HPLC) (Snyder et al., 2010) and gas chromatography–mass spectrometry (GC–MS) (Sparkman et al., 2011), provide accurate SCAAs measurements but require multi-step sample preparation, including protein hydrolysis and complex derivatization. They depend on specialized equipment, costly reagents, and long analysis times, which limit applicability for large-scale screening and rapid quality control in the food industry.

These limitations have motivated a shift toward spectroscopic methods that enable faster, more direct analysis. Vibrational spectroscopy methods, including infrared (IR), near-infrared (NIR), and Raman spectroscopy, provide detailed information on molecular structure, bonding, and composition (Bokobza, 1998; Ng & Simmons, 1999). IR-based techniques can be limited by water absorption and sample preparation requirements (Chon et al., 2021), whereas Raman spectroscopy is less affected by water and can be applied to aqueous extracts with minimal sample handling (Park et al., 2023). A practical limitation is that conventional Raman scattering is a low-probability phenomenon with weak intensity, reducing sensitivity for low-concentration analytes (Das & Agrawal, 2011). Surface-enhanced Raman spectroscopy (SERS) addresses this by using plasmonic nanostructures to amplify Raman signals and improve the sensitivity of detection for low-abundance analytes (Moskovits, 1985; Pilot et al., 2019). A defining property of quantitative SERS, in contrast to SERS for detection alone, is that under controlled experimental conditions, the scattered intensity is proportional to the number of molecules contributing to the enhancement. This means that spectral responses exhibit approximately proportional, monotonic behavior with analyte concentration, roughly analogous to Beer–Lambert–type relationships amenable to linear regression, and stable, repeatable patterns well-suited to machine learning (ML) analysis.

However, in complex food and biological matrices, SERS measurements are compromised by substrate heterogeneity, adsorption effects, fluorescence background, and non-linear baseline drift (Grys et al., 2021; Pilot et al., 2019). Conventional univariate or linear chemometric techniques often fail to decouple the target analyte signal from these complex, stochastic interferences. This limitation necessitates the use of artificial intelligence (AI) methods, including ML and deep learning (DL), to learn a quantitative mapping from SERS spectra to target analyte concentration. Recent label-free SERS studies show that ML/DL approaches can extract chemical and structural information from SERS spectra for discrimination and recognition tasks. Barucci et al. developed a hybrid strategy combining peak fitting with principal component analysis to discriminate proteins with closely similar spectral profiles, providing a reproducible approach to capture structure-dependent spectral variation in human and animal proteins (Barucci et al., 2021). Peng et al. implemented a DL-based, label-free SERS framework for screening and recognizing small-molecule binding sites in human drug-target proteins (Peng et al., 2022). While these studies focus on animal and human proteins and primarily address qualitative SERS analysis, related work in legume proteins supports extending this approach with ML/DL to quantitative analysis in complex food matrices (Findlay et al., 2025).

Accordingly, the present study develops and evaluates AI models to predict cysteine concentration from SERS spectra of pea (*Pisum sativum* L.) cultivars. Pea was selected as a representative legume matrix as pea protein is a widely used ingredient of growing importance in protein isolate production and processing, sustainable plant-based meat analogs and human nutrition (Shanthakumar et al., 2022). Peas are self-pollinating; thus, the pedigrees and lineages are well characterized, and cultivars exhibit genetic stability. Peas are the subject of well-developed breeding programs and cultivar collections, making them a good candidate for amino acid panel analysis by HPLC to establish reference values. Cysteine was selected as the analytical target because it contributes directly to the total SCAA pool and exhibits thiol-based surface-binding chemistry compatible with quantitative SERS (Findlay et al., 2025). A dataset of SERS spectra collected from 20 pea cultivars was used to investigate whether AI models can learn chemically meaningful relationships between spectral patterns and cysteine concentration.

To evaluate the complexity required to model these data, we selected five algorithms were selected, ranging from linear regression (LR) to convolutional neural networks (CNN). LR and partial least squares regression (PLSR) were included to establish a baseline and to represent standard chemometric approaches that assume linear spectral–concentration relationships. To assess whether non-linear modeling alone improves performance, we evaluated support vector regression (SVR) and random forest regression (RFR), which capture complex boundaries and variable interactions but rely on fixed input features. Finally, a DL model, a one-dimensional CNN (1D-CNN), was assessed. Unlike standard regression models that treat spectral points as independent features, CNNs are designed to learn hierarchical, local patterns such as peak shapes, widths, and relative shifts. This capability is hypothesized to make DL models more robust to the absolute intensity fluctuations and baseline shifts common in SERS, enabling better generalization across cultivars.

To assess this generalization capability, an evaluation framework was designed to distinguish between two forms of spectral variability that shape model behavior. The first, referred to as intra-cultivar spectral variability, arises from instrumental and substrate-related effects, including fluorescence background, stochastic noise, and local variations in electromagnetic field enhancement across the SERS substrate. These sources of variation occur within cultivars and reflect the physics of the measurement process. Therefore, they are assessed using a within-cultivar evaluation strategy, in which the training and test sets consist of spectra from the same cultivar (Section 2.3). The second form, termed inter-cultivar spectral variability, arises from cultivar-dependent biochemical variability, driven by genotype × environment (G×E) interactions that modify the molecular composition of pea extracts. This biochemical variation changes SERS peak intensities, peak positions, and baseline curvature across cultivars. To evaluate model performance under these conditions, we applied a leave-one-cultivar-out (LOCO) cross-validation strategy, which tests the model on an unseen cultivar excluded from the training process (Section 2.3). Evaluating AI models under both intra- and inter-cultivar spectral variability is critical for assessing their suitability for practical applications in legume breeding and quality assessment. For practical deployment in large-scale breeding programs or industrial quality control, analytical models must be able to predict analyte concentrations in new, unseen cultivars without requiring retraining. Consequently, the ability to generalize across genotypes, despite significant G×E biochemical variability, is a prerequisite for the operational utility of SERS-based screening.

Finally, we extended the evaluation to address two practical aspects of deployment: model interpretability and data acquisition efficiency. Shapley Additive Explanations (SHAP) were applied to identify the specific spectral features driving the predictions, ensuring that the model relies on chemically relevant vibrational modes. Separately, to optimize operational efficiency, a noise-modeling study was conducted to determine the minimum number of scans required for accurate prediction, providing practical guidelines for reducing acquisition time in high-throughput settings.

## 2. Materials and Methods

This section is divided into two main parts. Section 2.1 describes the experimental workflow and dataset generation, including the biological materials, sample preparation, SERS measurements, and the structure of the resulting spectral dataset. Section 2.2 details the AI analysis framework, including preprocessing, ML baselines, the DL architecture used to predict cysteine concentration from SERS spectra, and the model training and evaluation procedures. The overall workflow is summarized in Figure 1, which includes the post hoc model interpretability using SHAP and the noise-modeling study to guide scan count optimization.

**Figure 1.**
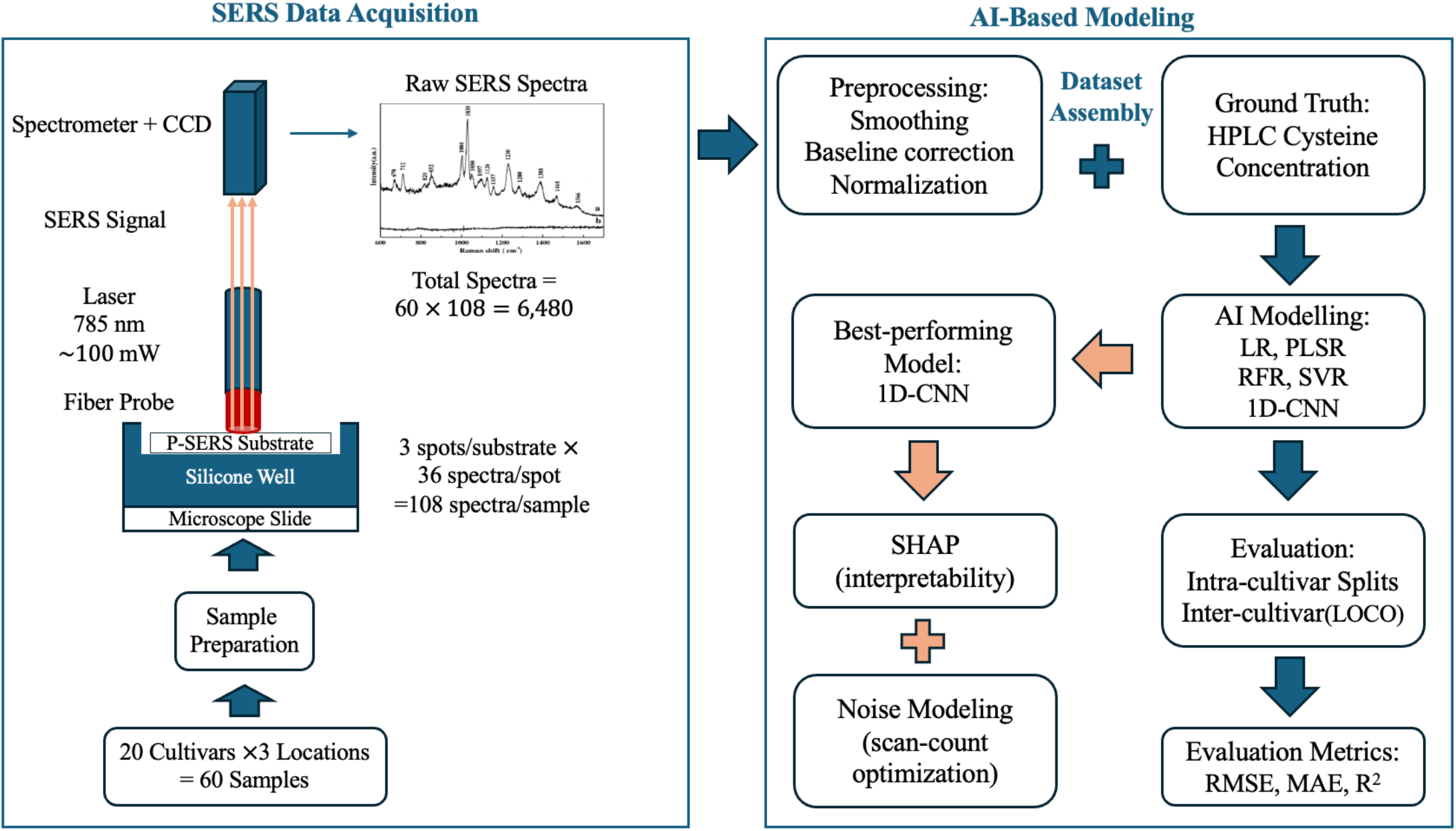
Overall workflow for SERS data acquisition and AI-based prediction of cysteine concentration. Left: SERS dataset generation from pea cultivars across three locations (60 samples). Sample extracts were prepared and measured on P-SERS substrates using a 785-nm excitation source and fiber-optic probe–based backscattering collection, with spectra acquired at multiple spots per substrate (3 spots/substrate, 36 spectra/spot; 108 spectra/sample; 6,480 total spectra). Right: AI-based modeling pipeline, including spectral preprocessing (smoothing, baseline correction, normalization) and dataset assembly by pairing preprocessed spectra with HPLC-derived cysteine concentrations (ground truth). Models included machine-learning baselines (LR, PLSR, SVR, RFR) and a 1D-CNN. Performance was evaluated using intra-cultivar splits and inter-cultivar (LOCO) testing, with RMSE, MAE, and R^2^ as evaluation metrics. The best-performing 1D-CNN was further analyzed using SHAP for interpretability and noise modeling to optimize scan count (acquisition time).

### 2.1. Overview of Experimental Data

This subsection describes the experimental workflow used to generate the SERS dataset for predicting cysteine concentration. The workflow includes cultivar selection, reference cysteine quantification, preparation of sample extracts, fabrication/preparation of SERS substrates, and SERS spectral acquisition. These steps produced a structured spectral dataset that serves as the input to the AI analyses described in Section 2.2.

#### 2.1.1. Pea Cultivars and Reference Compound

Flours from twenty pea cultivars from the CDC breeding program at the University of Saskatchewan, were selected based on their contrasting protein profiles to represent a diverse range: AAC Chrome, AAC Lacombe, AAC Liscard, CDC Amarillo, CDC Athabasca, CDC Canary, CDC Dakota, CDC Golden, CDC Greenwater, CDC Inca, CDC Jasper, CDC Striker, CDC Lewochko, CDC Meadow, CDC Patrick, CDC Saffron, CDC Spectrum, CDC Spruce, CDC Tetris, and Redbat 88. The prefixes in these names indicate their breeding origin: ‘AAC’ denotes varieties from Agriculture and Agri-Food Canada, and ‘CDC’ denotes those from the Crop Development Centre. Samples from each cultivar were ground into flour and analyzed for cysteine using the oxidative hydrolysis HPLC method described below. The corresponding HPLC reference cysteine concentrations for each cultivar across the three locations are provided in Table S1 (Supplementary Material).

Reference cysteine concentrations were determined using the performic acid oxidation–acid hydrolysis HPLC method described in (Findlay et al., 2025). In this procedure, proteins in pea flour extracts were oxidized with performic acid to convert cysteine (including disulfide-linked forms) to stable cysteic acid. Oxidized samples were then subjected to acid hydrolysis, derivatized using the AccQ-Tag Ultra reagent system, and separated on an AccQ-Tag Ultra C18 (1.7 μm) reversed-phase column using a Shimadzu UPLC system equipped with an SIL-30AC autosampler. Quantification was performed using calibration standards and hydrated amino acid molecular weights to obtain accurate cysteine-equivalent concentrations. L-cysteine (≥97%, Sigma-Aldrich) was used as the reference amino acid standard for calibration during HPLC analysis.

#### 2.1.2. Sample and Substrate Preparation

Alkaline extracts were prepared from pea flour samples (Section 2.1.1), and SERS spectral acquisition was performed following the method described by (Findlay et al., 2025) with modifications to reagent ratios and parameters described below. Pea flour was dispersed in Milli-Q water (0.2 g/1 mL). The suspension was kept on an ice bath and homogenized at a speed setting of 5 (∼20,000 rpm) for 30 seconds with an IKA Ultra-Turrax homogenizer (IKA-Werke GmbH & Co. KG, Staufen, Germany). Alkaline extraction was performed by adding 25 µL of a preprepared 1 M NaOH solution to 1.0 mL of pea homogenate, yielding a final pH of approximately 9. The mixture was vortexed for 10 seconds and incubated at room temperature (25 °C) for 2 hours. Samples were then centrifuged at 8000 rpm (∼5000 × g, rotor radius 7 cm) for 15 minutes at 4 °C. The supernatant was collected as the alkaline extract and stored at −80 °C.

Just prior to spectral acquisition, frozen extracts were thawed at room temperature and vortexed to ensure homogeneity. For each sample, 200 µL of extract was transferred into an individual silicone well and mixed with 100 µL of 20 mM tris (2-carboxyethyl) phosphine (TCEP) solution at pH 7, resulting in a final working volume of 300 µL. TCEP was added to reduce disulfide bonds and liberate free thiol for chemisorption to the SERS substrate. To prepare the SERS substrates, prefabricated paper-based SERS (P-SERS) substrates (Metrohm) were handled. Each substrate was positioned over a silicone well, and the handle was removed to allow the plasmonic surface tip to fall into the extract–TCEP mixture with the active side facing upward. The substrate was fully immersed and incubated at room temperature for 45 minutes to ensure consistent analyte–surface interaction. Following incubation, the substrates were transferred directly to the Raman system stage for spectral acquisition without drying or additional processing.

#### 2.1.3. SERS Spectral Acquisition

SERS measurements were acquired using a Raman system equipped with a 785 nm excitation laser delivering 100 mW at the sample surface. The system consisted of a Raman spectrometer coupled to a microscope-mounted sampling stage, enabling reproducible positioning of the P-SERS substrates beneath the Raman probe. Spectra were collected with a 1000 ms integration time and two co-additions, which were automatically combined by the instrument control software into a single stored spectrum, thereby balancing signal quality and measurement speed. Following the 45-minute incubation described in Section 2.1.2, each silicone well with P-SERS substrate was transferred directly to the Raman sampling stage while still immersed. The laser spot was focused on the submerged surface of the SERS substrate with the plasmonic sensing region oriented upward to maintain consistent optical alignment between the Raman probe and the active substrate surface.

To characterize nanoscale surface heterogeneity, spectra were acquired from three distinct spots on each P-SERS substrate. At each spot, 36 sequential spectra were collected without adjusting the optical focus or repositioning the substrate, yielding 108 spectra per sample. The same measurement protocol was repeated independently for samples obtained from each of the three Saskatchewan growing locations (Limerick, Rosthern, and Sutherland), yielding a total of 324 spectra per cultivar.

#### 2.1.4. Dataset

The complete dataset consisted of 6,480 raw SERS spectra (20 cultivars × 3 locations × 108 spectra). Each spectrum was a fixed-length vector of SERS intensities, indexed by Raman shift (cm^−1^), and was associated with a reference cysteine concentration determined by HPLC. All spectra from a given cultivar and location shared the same reference value. This yields a diverse dataset that captures both intra-cultivar spectral variability associated with instrumental and substrate-related effects (using P-SERS substrates from two separate manufacturing batches) and inter-cultivar spectral variability arising from cultivar- and location-dependent biochemical differences. It therefore supports a robust evaluation of model performance and generalizability for predicting HPLC cysteine concentration from SERS spectra.

### 2.2. AI-Based Modeling and Data Analysis Framework

This section describes the computational workflow used to develop models to predict HPLC cysteine concentration from SERS spectra. All computational steps were applied after the spectral dataset described in Section 2.1 was generated. The workflow includes spectral preprocessing, implementation of ML and DL models, and model evaluation.

#### 2.2.1. Spectral Preprocessing of SERS Data

Preprocessing is essential for preparing SERS spectra for AI-based modeling. Raw spectra often contain distortions, including baseline drift, random noise, fluorescence background, and intensity fluctuations, which can obscure chemically meaningful features and introduce non-chemical variability. Although many preprocessing algorithms exist, their suitability depends on the underlying physics of the spectroscopic technique. Accordingly, preprocessing should be tailored to the dominant sources of variability rather than applied in a generic manner.

In SERS and Raman spectra, one of the most prominent artifacts is the fluorescence background. It is a high-intensity, smoothly varying signal that can overwhelm Raman peaks and originates from both sample constituents and detector effects such as charge-coupled device (CCD) baseline drift (Bocklitz et al., 2011; Liland et al., 2016). SERS and Raman spectra are also affected by cosmic-ray artifacts, which appear as sharp, non-physical peaks resulting from high-energy particle impacts on the detector and can bias spectral analysis (Bocklitz et al., 2011; Wu & Chen, 2017). Additional distortions arise from Gaussian noise and other stochastic fluctuations inherent to Raman scattering, which reduce signal-to-noise ratios (S/N) in low-concentration measurements (Wahl et al., 2020). Fluctuations in laser power and changes in optical focusing further contribute to inconsistent peak heights across measurements. Another source of variation results from batch-to-batch differences in commercial SERS substrates. In this case, differences in nanostructure morphology and surface chemistry modify the local electromagnetic field distribution and change signal intensities across identical samples (Jeon et al., 2025). Numerous preprocessing methods have been proposed to address these issues, including baseline correction (Li et al., 2013; Lieber & Mahadevan-Jansen, 2003; Morháč & Matoušek, 2008; Peng et al., 2010), smoothing (Chen et al., 2013; Gorry, 1990; Kernel Smoothing - M.P. Wand, M.C. Jones), spike removal (Justusson, 1981; Li & Dai, 2011; Whitaker & Hayes, 2018), normalization, and derivative-based approaches (Chemometrics: Data Analysis for the Laboratory and Chemical Plant - Richard G. Brereton - Google Books, n.d.; Fearn et al., 2009).

In this study, SERS spectra were preprocessed using a workflow consisting of Savitzky–Golay (SG) smoothing (Savitzky & Golay, 1964), modified polynomial baseline correction (ModPoly) (Xia et al., 2018), and min–max normalization. First, SG smoothing was applied to reduce high-frequency noise while preserving peak shape. It applies a low-order polynomial filter within a moving window, where the window length and polynomial order control the degree of smoothing. Second, the ModPoly baseline correction was used to remove the fluorescence background and the slowly varying baseline curvature. This method estimates a polynomial baseline that captures the background trend of the spectrum, with the polynomial degree controlling its curvature. Cosmic-ray artifacts were addressed through the acquisition strategy rather than through post-processing. By collecting spectra with minimal co-addition, cosmic ray events were limited to single replicates rather than being averaged into the final spectra. This helps prevent attenuation of small but chemically relevant peaks. Finally, for the linear and kernel-based models (LR, PLSR, RFR, SVR), min–max normalization scaled each spectrum to the 0–1 range, thereby minimizing sensitivity to absolute intensity differences. In contrast, the 1D-CNN used unscaled spectral inputs, with internal scaling handled via batch-normalization layers (Section 2.2.2.5) rather than external normalization.

For each AI model described in Section 2.2.2, the preprocessing hyperparameters (SG window length, SG polynomial order, and ModPoly degree) were tuned using a grid search on the training set, with performance evaluated on a held-out validation set. The final preprocessing configuration used for each model is summarized in Table 1.

**Table 1.** Preprocessing configurations used for each model. All models use Savitzky–Golay (SG) smoothing and modified-polynomial baseline correction (ModPoly). LR, PLSR, RFR and SVR apply min–max normalization, whereas the 1D-CNN relies on internal normalization (batch-normalization layers).

| Model | SG Window Length | SG Polynomial Order | ModPoly Degree | Normalization |
| --- | --- | --- | --- | --- |
| LR | 11 | 3 | 3 | Min-Max |
| PLSR | 5 | 3 | 9 | Min-Max |
| RFR | 17 | 4 | 11 | Min-Max |
| SVR | 9 | 3 | 7 | Min-Max |
| 1D-CNN | 11 | 3 | 2 | - |

#### 2.2.2. Machine Learning and Deep Learning Models

To analyze the preprocessed SERS spectra, four ML algorithms were evaluated: LR, PLSR, RFR, and SVR. In addition, a 1D-CNN was used as the DL model. Each model was trained independently using the preprocessing workflow described in Section 2.2.1. The model descriptions, training procedures, and hyperparameter settings are outlined below. To ensure reproducibility, the complete source code and a sample dataset are openly available at https://github.com/Elhamm1/SERS-Data-Analysis/tree/main.

##### 2.2.2.1. Linear Regression

LR was used as a simple baseline model to describe the relationship between the preprocessed SERS spectra and cysteine concentration. For each spectrum ***x***, the predicted cysteine value *ŷ* was expressed as a linear combination of spectral intensities plus an intercept (*ŷ* = ***w***^***T***^***x*** + *b*), where ***x*** ∈ ℝ^*p*^ denotes the full preprocessed spectrum vector (with *p* = 1496 Raman shift bins), ***w*** denotes the regression coefficients and *b* is the intercept term. The parameters ***w*** and *b* were estimated by ordinary least squares, minimizing the sum of squared differences between predicted and HPLC-measured cysteine values in the training set, i.e., 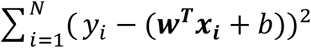. Because LR assumes a linear relationship between spectral features and cysteine concentration, it provides a baseline for comparison with more flexible ML and DL models.

##### 2.2.2.2. Partial Least Squares Regression

PLSR was included as a standard chemometric approach capable of handling strong collinearity across spectral variables. Unlike ordinary linear regression, which operates directly on the original intensities ***x***, PLSR projects each spectrum ***x*** ∈ ℝ^*p*^ into a lower-dimensional set of latent variables (components) that are constructed to maximize the covariance between the spectral features and cysteine concentration. The predicted cysteine concentration *ŷ* is then expressed as a linear combination of these latent variables plus an intercept, *ŷ* = *c*_1_*t*_1_ + *c*_2_*t*_2_ + ⋯ + *c*_*A*_*t*_*A*_ + *b*, where *t*_1_, …, *t*_*A*_ are the latent components extracted from the spectra ***x***, *c*_1_, …, *c*_*A*_ are the corresponding regression coefficients, *A* is the number of components, and *b* is an intercept term.

In this study, the optimal number of latent components was determined using five-fold cross-validation on a predefined search grid, yielding a final model with *A* = 25 components. The model was then refit using the selected number of components and used to generate predictions for the validation spectra. This procedure yields an LR model in a latent space aligned with cysteine variation and serves as a strong chemometric benchmark for comparison with the nonlinear ML and DL models.

##### 2.2.2.3. Support Vector Regression

SVR was used as a kernel-based nonlinear model to capture more complex relationships between the preprocessed SERS spectra and cysteine concentration. In SVR, the prediction for a new spectrum ***x*** is written as 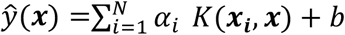, where ***x*** ∈ ℝ^*p*^, ***x***_*i*_ are training spectra, *α*_*i*_ are learned weights, *b* is an intercept term, and *K*(⋅,⋅) is a kernel function that defines the similarity between pairs of spectra and maps the data into a high-dimensional feature space.

In this study, a radial basis function (RBF) kernel was used to enable flexible, smooth nonlinear fits. The RBF kernel was defined as *K*(***x***_*i*_, ***x***_***j***_) = exp (−*γ* ∥ ***x***_*i*_ − ***x***_***j***_ ∥^2^), where ∥ ***x***_*i*_ − ***x***_***j***_ ∥ is the Euclidean distance between two spectra and *γ* controls the rate of decay of similarity with spectral distance. Model training was formulated as an optimization problem that keeps the regression function flat while constraining prediction errors within an *ε*-insensitive tube around the observed cysteine values. Deviations larger than *ε* are penalized through the regularization parameter *C*, which controls the trade-off between model complexity and error tolerance. Before SVR, input spectra were standardized to zero mean and unit variance. The hyperparameters *C, ε*, and *γ* were selected by five-fold cross-validation over a predefined grid using a Smooth L1 (Huber) loss, and the final SVR model used *C* = 5.0, *ε* = 0.01, and *γ* = 0.01. With this configuration, SVR provides a flexible nonlinear baseline that can model smooth spectral–concentration relationships while controlling model complexity through regularization and the kernel parameters.

##### 2.2.2.4. Random Forest Regression

Random Forest Regression was used to model nonlinear relationships between the preprocessed SERS spectra and cysteine concentration by combining many decision trees. The training data consist of pairs (***x***_*i*_, *y*_*i*_), where ***x***_*i*_ is the spectrum of the sample *i* (intensities at 1,496 Raman shift variables, denoted as p) and *y*_*i*_ is its HPLC-measured cysteine value. In this approach, a decision tree learns a sequence of binary split rules based on individual spectral variables. At each split, the algorithm selects a Raman shift variable and a threshold to reduce the variation in cysteine values across the resulting child nodes. This splitting process is repeated recursively and stops when further splits do not further reduce variation or when the remaining node contains few samples. The final nodes (leaf nodes) contain training spectra with similar cysteine values, and the prediction from a single tree for a new spectrum is the mean cysteine value of the training samples that fall into the same leaf. A random forest combines the predictions of many such trees to improve generalization and reduce variance.

In this study, *T* = 300 trees were trained. Each tree was fitted on a sample of the training data, and at each split, only a subset of spectral variables was considered as candidates, with the number of candidate features set to 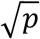. For a given spectrum ***x***, the forest prediction was obtained by averaging the outputs of all trees, 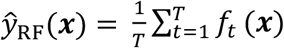, where *f*_*t*_(***x***) is the prediction from the *t*-th tree. This ensemble structure enables RFR to capture nonlinear dependencies and interaction effects among spectral features while reducing variance by averaging across multiple diverse trees.

##### 2.2.2.5. One-dimensional Convolutional Neural Network

1D-CNN was used as a DL model to capture nonlinear relationships between the SERS spectra and cysteine concentration. For an input spectrum **x** ∈ ℝ^1496^, the network treats **x** as a one-dimensional sequence with a single input channel and outputs a scalar prediction *ŷ* = *f*_*θ*_(**x**). For the *i*-th spectrum **x**_*i*_, this is *ŷ*_*i*_ = *f*_*θ*_(**x**_*i*_), where *f*_*θ*_ denotes the network parameterized by *θ*.

The architecture comprised four consecutive convolutional blocks, followed by two fully connected layers. The convolutional blocks used 1D convolutions with a kernel size of 5 and increasing numbers of filters (16, 32, 64, and 128). Each block applied convolution, batch normalization, and a ReLU activation, followed by max-pooling with a pool size (and stride) of 2 to downsample the spectral axis. These pooling operations reduced the spectral length by an overall factor of 16. The resulting feature maps were flattened and passed to a fully connected layer with 128 units and ReLU activation, followed by dropout (rate = 0.3) to reduce overfitting. A final linear output layer with a single neuron produced the predicted cysteine value.

The network was trained using mini-batch gradient descent with a batch size of 32. The training objective was a Smooth L1 (Huber) loss with *β* = 0.02. Optimization was performed using AdamW with an initial learning rate of 1 × 10^−4^ and decoupled weight decay, together with a OneCycle learning rate schedule over 100 epochs. Mixed-precision training and gradient clipping were used to stabilize optimization. Model performance was monitored on a held-out validation set at the end of each epoch, and the parameter set *θ* corresponding to the lowest validation loss was retained as the final 1D-CNN model. This architecture enables the 1D-CNN to learn hierarchical spectral features and capture nonlinear relationships between SERS patterns and cysteine concentration, while maintaining strong generalization via regularization.

### 2.3. Model Evaluation Strategy

To compare the ML and DL models, we used two evaluation strategies. First, we performed a within-cultivar split: approximately 80% of the spectra from each cultivar were used for training, and the remaining 20% were reserved for testing. Because the training and test spectra were from the same cultivar, the model was evaluated under conditions in which the spectral distribution was familiar. This setting provides a controlled baseline for assessing performance in the presence of intra-cultivar spectral variability and for tuning preprocessing and model hyperparameters. Second, we used a leave-one-cultivar-out (LOCO) cross-validation protocol to evaluate generalization across cultivars. In each LOCO fold, one cultivar was withheld as an independent test set, and the models were trained on spectra from the remaining 19 cultivars. This procedure was repeated until each cultivar had served once as the held-out test set. LOCO therefore measures how well a model generalizes to spectra from an unseen cultivar, assessing robustness against inter-cultivar spectral variability.

Model performance under both evaluation strategies was quantified using standard regression metrics. The root mean squared error (RMSE) was defined as 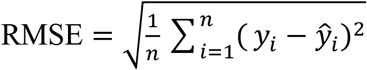, which places greater weight on larger errors. The mean absolute error (MAE) was calculated as 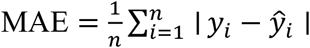, providing an average error measure in the same units as cysteine concentration, g/100 g. The coefficient of determination was computed as 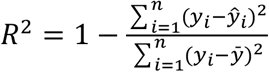, where *y* denotes the true cysteine concentration for the sample *i, ŷ* is the corresponding model prediction, 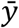 is the mean of the observed *y*_*i*_ values, and *n* is the number of spectra in the test set for a given split or LOCO fold. Together, these metrics summarize prediction performance under both within-cultivar and LOCO evaluations and enable a direct comparison of the ML models and the DL approach.

## 3. Results and Discussion

This section presents the results of the AI-based cysteine quantification and discusses their implications for model generalizability and practical deployment. As detailed in Section 2.1.4, the dataset comprised 6,480 SERS spectra collected across 20 pea cultivars. For the analysis, these spectra were stored in NumPy binary format, with each file containing a fixed-length vector of 1,496 Raman intensity values. All ML models were implemented using the scikit-learn library, while the 1D-CNN was implemented in PyTorch. Each model was trained on the same vectorized spectral inputs under the two evaluation strategies described in Section 2.3. These strategies were selected to distinguish between measurement-related artifacts (intra-cultivar) and biological diversity (inter-cultivar). Accordingly, Section 3.1 evaluates the predictive performance of the models under these different sources of spectral variability, while Section 3.2 extends beyond predictive performance to examine the interpretability and operational robustness of the 1D-CNN framework.

### 3.1. Impact of Spectral Variability on Model Performance

#### 3.1.1. Intra-Cultivar Spectral Variability

To assess model robustness against measurement noise, Table 2 summarizes the performance of the five models on raw versus preprocessed spectra. This comparison highlights how each algorithm responds to measurement-related artifacts, such as fluorescence background, baseline drift, and stochastic noise. For the four ML models (LR, PLSR, SVR, RFR), preprocessing yielded clear improvements, reducing RMSE and increasing R^2^. This suggests that these models are sensitive to baseline drift and noise, which can obscure the underlying spectral patterns. LR and PLSR benefited from baseline correction and scaling, which strengthened the linear relationship between spectral features and concentration. RFR also benefited from preprocessing, as reduced background and noise can yield more stable split decisions and more consistent averaging across trees. Finally, the improvement observed for SVR indicates that kernel-based similarity calculations are easily distorted by peak-shape noise.

**Table 2.** Performance of the five predictive models on raw and preprocessed SERS spectra. Results are reported as RMSE, MAE, and R^2^ for each model before and after applying Savitzky–Golay (SG) smoothing, modified polynomial baseline correction (ModPoly), and min–max normalization.

| Model | RMSE (g/100 g) | | MAE (g/100 g) | | $R^2$ | |
| --- | --- | --- | --- | --- | --- | --- |
|  | Raw | Preprocessed | Raw | Preprocessed | Raw | Preprocessed |
| LR | 0.013 | 0.012 | 0.010 | 0.008 | 0.561 | 0.650 |
| PLSR | 0.014 | 0.013 | 0.012 | 0.011 | 0.569 | 0.626 |
| RFR | 0.014 | 0.013 | 0.011 | 0.010 | 0.585 | 0.662 |
| SVR | 0.010 | 0.008 | 0.008 | 0.007 | 0.794 | 0.861 |
| 1D-CNN | 0.008 | 0.008 | 0.007 | 0.007 | 0.858 | 0.862 |

In contrast, the 1D-CNN achieved high performance (RMSE = 0.008 g/100 g, R^2^ = 0.862) on both raw and preprocessed inputs, showing no dependence on external preprocessing. This robustness arises from the convolutional layers, which process local spectral windows to capture the peak shape and structure rather than relying on absolute intensity. Additionally, internal mechanisms such as batch normalization and pooling effectively handle global intensity scaling and high-frequency fluctuations. Consequently, despite using substrates from two different manufacturing batches, the 1D-CNN can more effectively decouple the target signal from intra-cultivar measurement variability than other models.

#### 3.1.2. Inter-Cultivar Spectral Variability

To assess generalization to unseen genotypes, Table 3 compares model performance under within-cultivar and LOCO evaluation strategies. Under within-cultivar testing, where the test set contains familiar spectral distributions, all models performed reasonably well (RMSE 0.008–0.013 g/100 g). In this setting, variability is dominated by instrumental noise rather than by biochemical differences, allowing the models to fit the cultivar’s identity rather than its biochemical signature. However, under the LOCO evaluation, the ML models exhibited a significant performance decline when applied to unseen cultivars. R^2^ values dropped to 0.037–0.124, and RMSE increased by one order of magnitude. This indicates that these models rely on absolute peak intensities, which vary due to G×E interactions and substrate effects, rather than the intrinsic molecular signature of cysteine.

**Table 3.** Comparison of model performance under within-cultivar and LOCO evaluation schemes using preprocessed spectra.

| | RMSE (g/100 g) | | MAE (g/100 g) | | $R^2$ | |
| --- | --- | --- | --- | --- | --- | --- |
| Model | Within-cultivar | LOCO | Within-cultivar | LOCO | Within-cultivar | LOCO |
| LR | 0.012 | 0.097 | 0.008 | 0.045 | 0.650 | 0.037 |
| PLSR | 0.013 | 0.021 | 0.011 | 0.017 | 0.626 | 0.103 |
| RFR | 0.013 | 0.022 | 0.010 | 0.017 | 0.662 | 0.124 |
| SVR | 0.008 | 0.022 | 0.007 | 0.018 | 0.861 | 0.118 |
| 1D-CNN | 0.008 | 0.011 | 0.007 | 0.008 | 0.862 | 0.795 |

Conversely, the 1D-CNN demonstrated robust generalization, maintaining a low RMSE of 0.011 g/100 g and an R^2^ of 0.795 under LOCO conditions. This suggests that the convolutional architecture learns spectral features that are stable across cultivars. The network captures the local structure around each Raman peak and learns how intensities vary within small neighbourhoods, rather than relying on absolute peak heights, which vary across cultivars and substrates. It can learn detailed peak-shape characteristics, including curvature, width, and asymmetry, which are linked to molecular structure. These results confirm that while methods are sufficient for characterizing known samples, a 1D-CNN is required for generalizable prediction in breeding and quality-control applications, where new cultivars are encountered.

The validation of quantitative prediction and robustness to measurement variability addresses a key limitation highlighted in the Introduction. Prior work has used ML with SERS for qualitative objectives, such as discriminating protein types (Barucci et al., 2021) or identifying binding-related spectral changes (Peng et al., 2022). Here, we benchmark AI-based SERS analysis for quantitative regression in a complex food matrix using a deployment-focused evaluation. It is shown that the 1D-CNN maintains strong performance when applied to unseen cultivars. To the best of our knowledge, this is the first application of deep learning to quantify a specific amino acid (cysteine) in legume extracts using SERS. This advancement establishes a scalable framework for high-throughput phenotyping, enabling breeders to rapidly screen for nutritional quality.

### 3.2. Practical Applications of the 1D-CNN for Cysteine Quantification in Pea Cultivars

This section extends beyond predictive performance to examine how the 1D-CNN can be used to support SERS-based quantification of cysteine in this study. First, we use SHAP to identify the Raman regions that most contribute to model predictions under both evaluation schemes. Second, we evaluate the model’s sensitivity to spectral noise using a controlled augmentation framework that simulates varying scan counts and quantifies their effects on predictive performance.

#### 3.2.1. Interpreting Raman Vibrational Features Across Cultivars

To examine which Raman regions the 1D-CNN uses for cysteine prediction, we applied SHAP analysis to the trained 1D-CNN models under both evaluation schemes. For the within-cultivar split, SHAP values were computed for the corresponding within-cultivar model. For LOCO (20-fold), SHAP values were computed using the best-performing fold. Figure 2 shows SHAP summary plots for the within-cultivar model (left) and the LOCO model (right). Each point corresponds to a spectrum in the SHAP evaluation set. Features are ranked from top to bottom by mean absolute SHAP value, which reflects the average magnitude of the contribution of each feature to the predicted cysteine concentration across spectra. The horizontal axis shows SHAP values, indicating whether each feature increases or decreases the predicted cysteine concentration. Point color indicates the Raman intensity at that Raman shift, from low to high.

**Figure 2.**
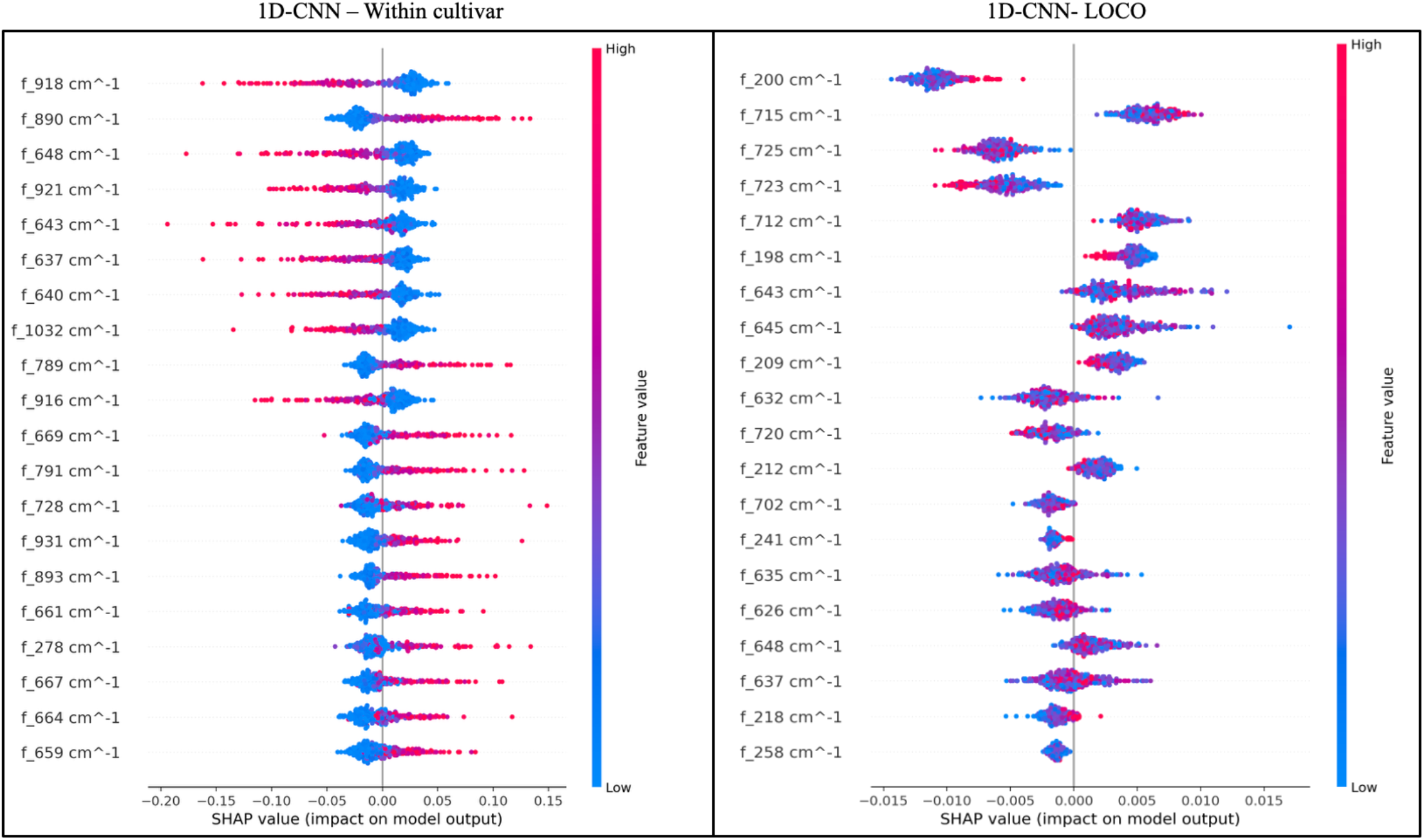
SHAP summary plots showing Raman-shift regions that contribute to 1D-CNN predictions of cysteine concentration under two evaluation schemes: within-cultivar split (left) and leave-one-cultivar-out (LOCO) evaluation (right). Features (Raman shift, cm^−1^) are ordered from top to bottom by mean absolute SHAP value, representing the average contribution magnitude to the model output. Each point corresponds to one spectrum in the SHAP evaluation set. The x-axis shows SHAP values (in the units of the model output), where positive values increase the predicted cysteine concentration and negative values decrease it. Point color indicates the feature value at that Raman shift, from low to high.

In the within-cultivar setting (Figure 2, left), the most impactful features are distributed across multiple Raman-shift regions rather than concentrated in a single band. Dominated contributions appear both near ∼880–930 cm^−1^ (e.g., 890, 918, 921, 931 cm^−1^) and within the ∼630–650 cm^−1^ region (e.g., 637, 640, 643, 648 cm^−1^), with additional contributions at intermediate bands (e.g., ∼669–791 cm^−1^). This pattern suggests that when cultivar identity is shared between training and testing, the model can rely on a broader set of spectral features rather than on a single feature associated with cysteine.

In the LOCO setting (Figure 2, right), the SHAP ranking is more structured across Raman regions. Although a low-Raman shift feature near ∼200 cm^−1^ appears as a top contributor, most of the highly ranked features are concentrated in the ∼630–760 cm^−1^ range (e.g., ∼632–648 cm^−1^ and ∼702–725 cm^−1^). Low-Raman-shift SERS features near ∼200 cm^−1^ are attributed to substrate-related contributions, such as metal–adsorbate interactions, metal lattice/phonon modes, or electronic scattering, rather than to internal molecular vibrations (Inagaki et al., 2019). In addition, Ag P-SERS spectra show a dominant low-shift band near ∼235 cm^−1^ that has been assigned to Ag–Ag stretching (Findlay et al., 2025), consistent with substrate-related contributions at low Raman shifts. In contrast, biochemical interpretation is supported by the highly ranked bands in the 630–760 cm^−1^ region (Adar et al., 2022). Features near ∼643–648 cm^−1^ and ∼712–725 cm^−1^ are consistent with reported protein carbon–sulfur (C–S)–related vibrations in this band range, supporting their relevance for cysteine prediction under LOCO. The LOCO ranking provides the most appropriate basis for interpreting 1D-CNN behavior in cross-cultivar prediction, because it reflects features that remain informative when the test cultivar is not represented in the training data.

#### 3.2.2. Data Acquisition Optimization and Noise Modeling

The controlled-noise study examines how a 1D-CNN can inform practical choices in SERS data acquisition. As described in Section 2.1.3, each stored spectrum was acquired with two co-additions, meaning that two consecutive acquisitions were combined into a single spectrum. In the noise-modeling analysis, the scan count *N* refers to the effective number of averaged acquisitions per spectrum (co-additions). Because random noise decreases with increasing *N*, spectra acquired with fewer effective scans have lower signal-to-noise ratios. Quantifying how prediction performance changes as the number of scans decreases is therefore important for balancing acquisition time against analytical performance. To assess this trade-off, we used a noise-modeling and augmentation framework to generate synthetic spectra that mimic measurements at different effective scan counts while preserving the underlying spectral structure of the original data.

The augmentation strategy was based on signal-averaging theory, where the standard deviation of random noise scales as 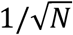. We defined a high-SNR reference level *N*_ref_ = 512, and scaled the additive noise by 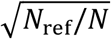 to simulate effective scan counts from 64 down to 1. The reference level *N*_ref_ = 512 was chosen to provide a wide signal-to-noise range to resolve performance trends across scan counts. It is used only as a reference for scaling the added noise and does not imply that spectra were experimentally acquired with 512 co-additions. This approach generated datasets with increasing noise while preserving the underlying spectral structure.

To perform the noise augmentation, we randomly selected 10 spectra from the 324 spectra available for each cultivar and generated augmented versions of these spectra for each simulated scan count. The results, summarized in Table 4, show a relationship between scan count and predictive performance. As the number of scans decreases from 64 to 1, RMSE increases from 0.009 to 0.016, MAE increases from 0.008 to 0.014, and R^2^ decreases from 0.843 to 0.446. Improvements in model performance become smaller beyond 16 scans. The model maintains low RMSE and higher R^2^ at 64, 32, 16, and 8 scans. At 4 and 2 scans, RMSE increases to 0.014 and R^2^ falls to 0.607 and 0.583. Performance is lowest at 1 scan. These results indicate that the 1D-CNN is robust to noise as the number of scans is reduced. Based on these results, 8 scans provide a good balance between acquisition time and predictive performance. Using 2 co-additions, consistent with the experimental protocol (Section 2.1.3), is also possible but yields lower accuracy than 8 scans.

**Table 4.** Performance of the 1D-CNN model as a function of simulated scan count in the noise-modeling experiment. Spectra at each scan level were generated by scaling additive noise relative to a 512-scan reference. Performance is reported as RMSE, MAE, and R^2^ for cysteine concentration.

| Number of Scans | RMSE | MAE | $R^2$ |
| --- | --- | --- | --- |
| 64 | 0.009 | 0.008 | 0.843 |
| 32 | 0.010 | 0.009 | 0.777 |
| 16 | 0.010 | 0.009 | 0.774 |
| 8 | 0.011 | 0.009 | 0.770 |
| 4 | 0.014 | 0.012 | 0.607 |
| 2 | 0.014 | 0.013 | 0.583 |
| 1 | 0.016 | 0.014 | 0.446 |

Beyond evaluating noise effects, this analysis highlights the application of data augmentation when experimental data are limited. By generating realistic spectral variants that expand the training set, the augmentation procedure allows the 1D-CNN to learn a broader range of instrumental noise, baseline variation, and spectral fluctuations. This is particularly useful in SERS studies, where data collection is often constrained by sample availability, instrument time, or substrate variability.

## 4. Conclusion

This study demonstrates that AI-based modeling of SERS spectra enables the quantitative prediction of cysteine in pea extracts. Model performance depended on the dominant source of variability represented in the evaluation. Under within-cultivar testing, where intra-cultivar spectral variability dominates, the ML models benefited from preprocessing and achieved moderate-to-high performance. In contrast, when the evaluation introduced inter-cultivar spectral variability through LOCO testing, the performance of traditional regression models declined sharply, indicating weak generalization to unseen cultivars. The 1D-CNN showed better cross-cultivar generalization, with only a small increase in RMSE from within-cultivar to LOCO testing, supporting its suitability for applications where new cultivars are expected at deployment.

SHAP analysis provided insight into how the 1D-CNN interprets behaves under intra- and inter-cultivar spectral variability. Within the cultivar, feature importance was distributed across multiple regions. Under LOCO conditions, feature importance became more structured and concentrated in the ∼630–760 cm^−1^ region, with an additional contribution from a low-Raman shift feature near ∼200 cm^−1^. The concentration of influential features in the 630–760 cm^−1^ range, which is consistent with reported C–S–related vibrational contributions in proteins, supports a chemical basis for cross-cultivar prediction and confirms the identification of spectral patterns that remain stable across cultivars. From a practical perspective, the noise study indicates that with 8 scans, the 1D-CNN maintained performance comparable to that obtained with 16 or 32 scans, thereby reducing acquisition time. In addition, the model demonstrated consistency across batches with respect to substrate variability, thereby addressing a barrier to SERS reproducibility. It confirms its suitability for routine operations where consumable properties vary.

Overall, the findings support the use of SERS combined with DL as a practical and scalable approach for rapid, cross-cultivar prediction of cysteine concentration, thereby supporting food-quality control and cultivar selection. Future work should extend the approach to full-panel amino acid profiling and compare the current 1D-CNN with alternative DL architectures to evaluate improvements in generalization and robustness across diverse plant protein matrices.

## Acknowledgements

This work was supported by the National Research Council of Canada (NRC) through the Sustainable Protein Production (SPP) Program grant number SPP-142–1. The authors also acknowledge Obasi Ukpai Ukoji and Sristi Mundhada for their contributions to the SERS data acquisition.

