## Supplemental Table 1 for "Generalizable Cysteine Quantification in Pea Cultivars from SERS Spectra Using AI"

**Table S1.** Mean cysteine concentration (g/100 g) across three field locations (Limerick, Rosthern, Sutherland) for 20 pea cultivars, with the corresponding standard deviation across locations.

| # | Cultivar | Mean (g/100g) | SD (g/100g) |
|---|---|---|---|
| 1 | AAC chrome | 0.317211 | 0.012077 |
| 2 | AAC Lacombe | 0.338561 | 0.029327 |
| 3 | AAC Liscard | 0.325405 | 0.044341 |
| 4 | CDC Amarillo | 0.358930 | 0.053315 |
| 5 | CDC Athabasca | 0.341203 | 0.015273 |
| 6 | CDC Canary | 0.314476 | 0.007967 |
| 7 | CDC Dakota | 0.332835 | 0.031283 |
| 8 | CDC Golden | 0.359066 | 0.031501 |
| 9 | CDC Greenwater | 0.346381 | 0.006835 |
| 10 | CDC Inca | 0.365342 | 0.023049 |
| 11 | CDC Jasper | 0.337341 | 0.007358 |
| 12 | CDC Striker | 0.344968 | 0.025169 |
| 13 | CDC Lewochko | 0.341267 | 0.039327 |
| 14 | CDC Meadow | 0.312012 | 0.012532 |
| 15 | CDC Patrick | 0.338404 | 0.053002 |
| 16 | CDC Saffron | 0.346789 | 0.026855 |
| 17 | CDC Spectrum | 0.373175 | 0.042144 |
| 18 | CDC Spruce | 0.345676 | 0.013716 |
| 19 | CDC Tetris | 0.342040 | 0.019235 |
| 20 | Redbat 88 | 0.316257 | 0.023543 |
